# Single-Molecule FRET at 10 MHz Count Rates

**DOI:** 10.1101/2023.12.08.570755

**Authors:** Lennart Grabenhorst, Flurin Sturzenegger, Moa Hasler, Benjamin Schuler, Philip Tinnefeld

## Abstract

A bottleneck in many studies utilizing single-molecule Förster Resonance Energy Transfer (smFRET) is the attainable photon count rate as it determines the temporal resolution of the experiment. As many biologically relevant processes occur on timescales that are hardly accessible with currently achievable photon count rates, there has been considerable effort to find strategies to increase the stability and brightness of fluorescent dyes. Here, we use DNA nanoantennas to drastically increase the achievable photon count rates and to observe fast biomolecular dynamics in the small volume between two plasmonic nanoparticles. As a proof of concept, we observe the coupled folding and binding of two intrinsically disordered proteins which form transient encounter complexes with lifetimes on the order of 100 μs. To test the limits of our approach, we also investigated the hybridization of a short single-stranded DNA to its complementary counterpart, revealing a transition path time of 17 μs at photon count rates of around 10 MHz, which is an order-of-magnitude improvement when compared to the state of the art. Concomitantly, the photostability was increased, enabling many seconds long megahertz fluorescence time traces. Due to the modular nature of the DNA origami method, this platform can be adapted to a broad range of biomolecules, providing a promising approach to study previously unobservable ultrafast biophysical processes.

## Introduction

Single-molecule Förster Resonance Energy Transfer (smFRET) experiments are a cornerstone in the investigation of biomolecular dynamics [1] as they enable the observation of the molecule of interest ‘in action’, specifically within its natural environment, and give a real-time movie that can span timescales from milliseconds to minutes or in some cases even hours [2]. Crucially, although methods enabling access to faster timescales through correlation techniques exist [3, 4] time-resolving individual rare events, such as the rapid jumps across the barriers between two conformational states (e.g. protein or nucleic acid folding), requires very high count rates from single molecules [5, 6]. These transition paths have gained increasing interest and smFRET experiments have been playing a major role for revealing the nature and timescales of these paths [7, 8]. However, the photon count rates (PCR) required for accessing the relevant microsecond timescale mandate the use of high excitation intensities leading to saturation and increased photobleaching, which makes the task of measuring these transition paths very challenging. Plasmonic hotspots have been shown to increase the photostability and brightness of fluorescent labels [9, 10], and offer a potentially complementary strategy to the more commonly used chemical photostabilization [11–13]. Coupling the emitters to plasmonic hotspots not only increases the electric field between the two plasmonic nanoparticles, but also increases the emission rates of the fluorophore, leading to a shorter time that the molecule spends in the reactive excited states as well as reducing the time until the fluorophore is available for re-excitation [14, 15]. This, in turn, results in improved photostability of fluorescent labels and also enables higher fluorescence intensities without saturation [9, 16, 17]. First examples employing similar strategies for diffusing molecules have appeared and showed promising results [18, 19], but they are limited to the submillisecond timescales of molecular diffusion through the excitation volume, which makes the complete observation of complex biomolecular pathways unlikely. DNA origami nanoantennas [20, 21] have overcome the challenge of selective immobilization of biomolecules in plasmonic hotspots. Recent iterations have optimized the usability of the plasmonic hotspots in NACHOS (NAnoantennas with Cleared HOtSpots) improving binding kinetics and the immobilization of larger functional entities such as diagnostic assays [22] and even antibodies [23] in these zeptoliter volumes. In this work, we explore the use of NACHOS for biophysical single-molecule FRET experiments. Recent studies have shown that the electromagnetic environment of plasmonic hotspots has an influence on the FRET rate coefficient as well as the FRET efficiency [24–26], but their use in biophysics is largely unexplored.

Here, we optimize DNA origami nanoantennas for biophysical experiments with a careful selection of parameters, including the choice of donor and acceptor fluorophores. With optimized hotspots, we demonstrate strongly enhanced countrates and long-lasting single-molecule FRET time traces. Unaltered biomolecular function is demonstrated by reproducing the lifetime of a short-lived intermediate in the coupled folding and binding of two intrinsically disordered proteins. Transition path times of down to a few microseconds are then reported for a DNA hybridization reaction.

## Results and Discussion

An important first step of this endeavor is the choice of a suitable FRET pair. Photophysical characteristics such as the formation of short-lived dark or dim states [9] can complicate the analysis of the data and need to be avoided or accounted for in the analysis. Furthermore, both fluorophores should be spectrally red-shifted with respect to the plasmon resonance of the gold dimer nanoantenna [27]. In consideration of these aspects, we examined FRET pairs covering the red and near IR spectral region (a deeper discussion on dye selection and DNA origami nanoantenna production is found in Supplementary Note 1).

First, we investigated the binding of the nuclear-coactivator binding domain of the CBP/p300 transcription factor (NCBD) to the activation domain of SRC-3 (ACTR). Previous studies with smFRET [28] revealed that these intrinsically disordered proteins (IDPs) form a transient encounter complex before they form a fully folded heterodimer. The lifetime of this encounter complex has been determined experimentally to approximately 80 μs – 100 μs, depending on the ionic strength of the buffer [28], which offers a way to test whether our system is compatible with measuring protein dynamics. Using a single-stranded DNA (ssDNA) handle attached to a cysteine residue close to the N-terminus, we immobilized ACTR labelled with AlexaFluor 647 in the plasmonic hotspot (Fig. 1a, b). Free NCBD labelled with LD750 was then added to the imaging buffer which led to the appearance of short segments of high FRET efficiency in the fluorescence recordings, indicating the binding of NCBD to ACTR. To identify nanoantennas showing reversible binding events without prematurely photobleaching the FRET pair, we started the acquisition of single-molecule fluorescence trajecories at lower excitation intensities (10 – 20 nW). Then, we increased the excitation intensity (1 – 4 μW) to acquire high intensity time traces. With the fluorescence enhancement and increased photostability provided by the DNA origami nanoantenna, we were able to increase the average photon count rate by almost 3-fold to 588 kHz compared to earlier studies with a different FRET pair [28] (see Supplementary Note 2) and achieved a maximum photon count rate of over 2 MHz, with a mean observation time of 0.66 s, which also represents a significant improvement. While intermediates with longer lifetimes can sometimes be directly observed in the time traces (Fig. 1c, left), quantifying intermediates with shorter lifetimes remains challenging (Fig. 1c, right, see Fig. S1 for more examples). Consistent with previous mechanistic studies of FRET in plasmonic nanostructures [24–26], the FRET efficiency of the bound state is decreased in the plasmonic hotspot — we obtained a mean value of ⟨*E*_B_⟩ = 0.49 (uncorrected, compare Fig. S7)

**Figure 1:**
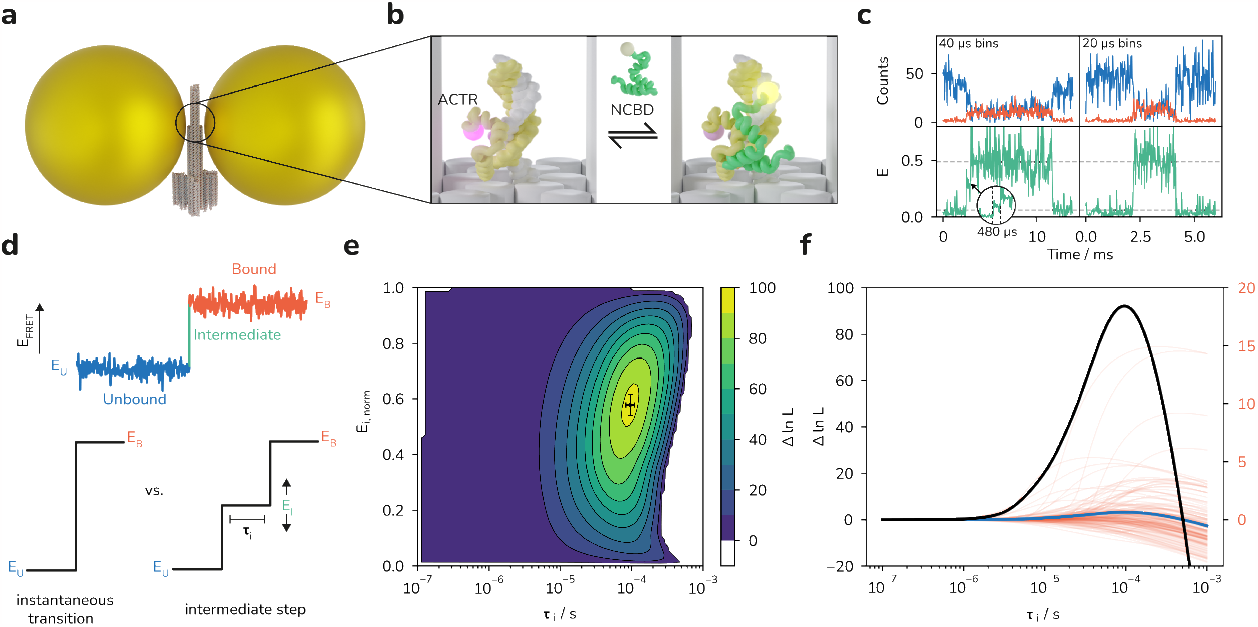
Investigation of the coupled folding and binding of ACTR to NCBD using DNA origami nanoantennas. **(a)** Illustration of the DNA origami nanoantenna with 100 nm gold nanoparticles. **(b)** Depiction of the binding reaction of ACTR to NCBD. We used ssDNA handles to immobilize ACTR (yellow) labelled with AlexaFluor 647 in the plasmonic hotspot region. NCBD (green) labelled with LD750 is freely diffusing in solution and transiently binds to ACTR. Cartoon representations were obtained from the PDB [29, 30]. **(c)** Exemplary fluorescence time traces (orange: acceptor, blue: donor) and corresponding uncorrected FRET efficiencies (E, green) of binding events represented with 40 μs (left) and 20 μs (right) binning. The inset on the left highlights a potential intermediate state. The dashed lines indicate the mean values for bound (*E*_B_) and unbound (*E*_U_). **(d)** Schematic depiction of the two models that are compared in the maximum likelihood analysis [7]: an instantaneous transition from the unbound to the bound state is compared to a transition that includes an intermediate step with FRET efficiency *E*_I_ and of lifetime τ_i_. **(e)** 2D contour plot of the log-likelihood difference ∆ ln *L* between the two models summed up over 141 transitions. The most likely values are τ_i_ = 96 ± 13 μs and *E*_i,norm_ = 0.58 ± 0.04. The errors were calculated from the diagonal elements of the covariance matrix. **(f)** Plot of ∆ ln *L* values versus τ_i_ for single transitions (red, right scale) and the sum of all transitions (black) as well as their average (blue, right scale) calculated at the most likely value for *E*_i,norm_.

In order to fully quantify the most likely lifetime of the intermediate state, we used the photon-by- photon maximum likelihood approach [7, 28, 31]. Here, the likelihood of a transition between two states via an intermediate with FRET efficiency *E*_I_ and lifetime τ_i_ is compared to the likelihood of an instantaneous transition (Fig. 1d). To account for acceptor blinking, we included an additional dark state in the analysis [32]. We also normalized each *E*_I_ value to the respective transition-wise bound state (*E*_B_) and unbound state (*E*_U_) FRET efficiency values: *E*_i,norm_ = (*E*_I_-*E*_U_)/(*E*_B_-*E*_U_), to account for the slightly broadened FRET distributions resulting from the heterogenous fluorescence enhancement by the nanoantenna (see SI for details on the analysis). This means that for an *E*_i,norm_ value of 0.5, the intermediate state lies exactly halfway between *E*_U_ and *E*_B_. As we noticed a decrease of the total photon count rate of the system upon binding of NCBD (Fig. S1), we additionally carried out simulations of binding events with non-constant total photon count rates and found no significant influence on the robustness of the analysis (see Supplementary Note 3 and Figs. S3 and S4). From a total of 141 transitions, we obtained a value of 96 ± 13 μs for the most likely lifetime of the intermediate state and an *E*_i,norm_ value of 0.58 ± 0.04 (Fig. 1e–f). These values are in excellent agreement with previously reported values at similar ionic strength (90 ± 10 μs [28]), which illustrates the potential of our approach to be applied to other biomolecular processes.

To test our approach on even faster reactions and simultaneously avoid potential negative effects on the photostability of our labels by the neighboring amino acids and labelling chemistries (e.g. cysteine or maleimide [33]), we next set out to investigate the transient binding of a short ssDNA to a ssDNA docking site placed in the plasmonic hotspot. DNA-DNA hybridization is believed to occur via a 2– 3 nucleotide (nt) nucleation site which then transforms into the stably bound state via zippering of the remaining bases [34]. In our experiments, we used a 6-nt long ssDNA labelled with Dy-751 at the 3’-end, which hybridizes to a docking site labelled at the 3’-end with a Cy5B [35, 36] fluorophore (Fig. 2a). We achieved a mean PCR of 2.42 MHz and a maximum count rate of 8.93 MHz, which is approximately an order of magnitude higher than previously reported PCRs in smFRET experiments [7, 37, 38] and also substantially higher than what we observed in the experiments described earlier. It is worth emphasizing that because of the increased photostability, we were sometimes able to observe the same molecule for up to 30 seconds at photon count rates of 2 MHz and record up to 57 million photons from one FRET pair (Fig. 2b). This allowed us to occasionally observe more than 20 transitions on one molecule, which illustrates how the nanoantenna approach facilitates data acquisition. Figure 2c shows two exemplary snapshots from fluorescence time traces of a binding event (see Fig. S2 for more examples). At the chosen 2–4 μs bin sizes, acceptor blinking becomes visible (indicated by arrows), substantiating the need to account for it in the analysis. We observed a total of 405 transitions and analyzed these according to the photon-by-photon maximum likelihood approach described above. We found a mean value of the uncorrected FRET efficiency of the bound state of 0.48, a most likely transition path time of 17 ± 1 μs and an *E*_i,norm_ value of 0.30 ± 0.02 (Fig 2b). To test the influence of the FRET pair on the results, we repeated the experiment with AlexaFluor 647 as donor and ATTO740 as acceptor fluorophores. Although we were able to achieve slightly higher count rates (mean count rate: 5.34 MHz, with maximum count rates of up to 16.4 MHz, see Fig. S3), the FRET efficiency of the bound state was notably lower (⟨ *E*_B_⟩ = 0.30), potentially making it more difficult to distinguish between the bound and the unbound states. This can at least partially be explained by the lower Förster radius of this dye pair (see Table S3). Nevertheless, we still obtained a transition path time that agrees with the one measured with the Cy5B/Dy-751 FRET pair (τ_i_ = 20 ± 3 μs) with a most likely value for *E*_i,norm_ of 0.21 ± 0.02.

**Figure 2:**
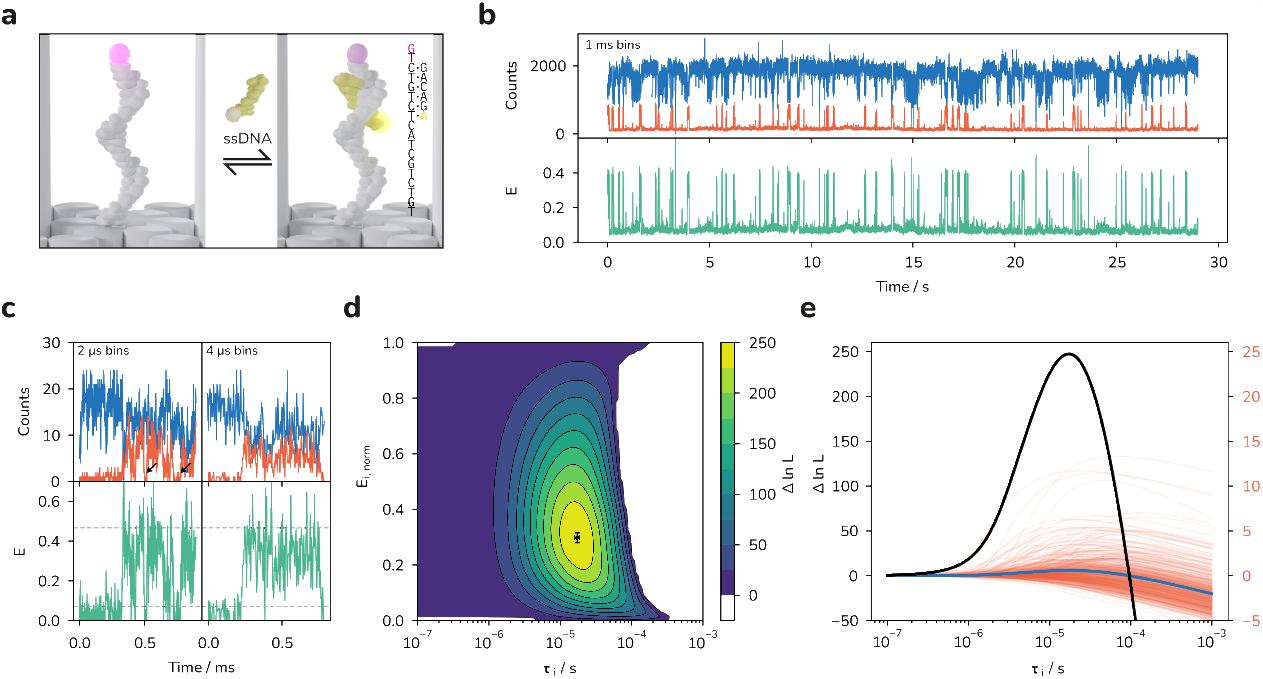
Investigation of DNA-DNA hybridization reactions in the plasmonic hotspot. **(a)** Illustration of the experiment: a 6 nt ssDNA labelled with Dy-751 hybridizes to a ssDNA docking site labelled at the 3’- end with a Cy5B fluorophore. The DNA sequences are shown in the inset. **(b)** The increased photostability allows for the observation of several binding events at high count rates. Top: acceptor (orange) and donor (blue) fluorescence time trace at 1 ms binning. Bottom: corresponding uncorrected FRET time trace. **(c)** Exemplary fluorescence time traces (top) showing donor (blue) and acceptor (orange) fluorescence during a hybridization event at 2 and 4 μs binning. The corresponding uncorrected FRET time trace is shown below. The arrows indicate possible acceptor blinking events. **(d)** 2D contour plot of the log likelihood difference ∆ ln *L* of a transition of duration τ_i_ with FRET efficiency Ei, norm to an instantaneous transition summed over 405 transitions. The most likely values are τ_i_ = 17 ± 1 μs and *E*_i,norm_ = 0.30 ± 0.02. The errors are calculated from the diagonal elements of the covariance matrix. **(e)** Plot of ∆ ln *L* values versus τ_i_ for single transitions (red, right scale) and the sum of all transitions (black) as well as their average (blue, right scale) calculated at the most likely value for *E*_i,norm_.

Previous FRET studies on DNA hairpin formation were able to determine an upper bound of 2.5 μs for the hybridization of a 4 nt long hairpin stem [39] and optical tweezer experiments reported values around 30 μs for much longer hairpins [40]. Our results thus lie in a plausible range of transition path times for 6-nt DNA-DNA hybridization event, although further experiments will be needed to fully understand the process of hybridization. For example, in preliminary experiments, the inclusion of an AT-mismatch in the short strand labelled with ATTO740 led to an increase of τ_i_ to 41 ± 3 μs with a similar value for *E*_i,norm_ of 0.229 ± 0.012 (see Fig. S4).

## Conclusion

In summary, we showed that DNA origami nanoantennas can be used to accelerate data acquisition and increase the attainable photon count rates in smFRET experiments on biomolecular dynamics. As a proof of concept, we investigated the coupled folding and binding of two IDPs as well as the hybridization of two ssDNAs and showed that we could resolve processes on the microsecond timescale. With the resulting higher time resolution [41, 42], previously inaccessible biological processes could be investigated. To this end, the modular nature of the DNA origami technique allows the swift incorporation of new biomolecules of interest, making this platform broadly applicable as a tool for the improvement of temporal resolution in smFRET experiments. Another benefit which we did not focus on in this work is the possibility to work at much higher concentrations of labelled species due to the ultra-small volume of fluorescence enhancement in the nanoantennas, which also allows the observation of biomolecular interactions with micromolar dissociation constants [43] – we estimated the dissociation constant of the mismatched DNA-DNA interaction (Fig. S4) to 200 μM and still were able to clearly resolve the binding events at 3.6 μM concentration of the short strand in solution (Fig. S8). Recent iterations of the DNA origami nanoantenna could accommodate even larger biomolecules and should further reduce the potential impact of the constricted environment on the experiments [44]. The development of new near-infrared fluorophores showing less blinking on the low microsecond timescale [45] which potentially interferes with the maximum-likelihood analysis should make it possible to further push the time resolution and possibly, in combination with more efficient sampling methods, bridge the gap towards the timescales of molecular dynamics simulations [46–48].

## Acknowledgements

L.G. thanks Viktorija Glembockyte for many insightful discussions. The authors thank Martin J. Schnermann and Ryan McLaughlin for providing the Cy5B dye. F.S., M.H., and B.S. thank Daniel Nettels for discussion and providing data analysis tools. This work was supported by the German research foundation (DFG, grant number TI 329/9-2, project number 267681426, INST 86/1904-1 FUGG, excellence cluster e-conversion EXC 2089/390776260), the Free State of Bavaria under the Excellence Strategy of the Federal Government and the Länder through the ONE MUNICH Project Munich Multiscale Biofabrication, and the Swiss National Science Foundation (B.S.).

## Notes

### Competing Interest Statement

The authors have declared no competing interest.

