## Supplementary Information for "Single-Molecule FRET at 10 MHz Count Rates"

**Supporting information**

Lennart Grabenhorst[1], Flurin Sturzenegger[2], Moa Hasler[2], Benjamin Schuler[2,3], and Philip Tinnefeld[1,*]

[1]Department of Chemistry and Center for NanoScience, Ludwig-Maximilians-Universität München, München, Germany
[2]Department of Biochemistry, University of Zurich, Zurich, Switzerland
[3]Department of Physics, University of Zurich, Zurich, Switzerland

**3 Supplementary tables** **9**

**4 Supplementary figures** **11**

**References** **17**

### 1 Materials and methods

#### 1.1 DNA origami synthesis

DNA origami nanostructures were produced and purified as described previously. [1] Purification of the DNA origami was achieved using 100 kDa MWCO filters (Merck). The filters were filled with 450 µL FoB5 buffer (10 mM TRIS, 5 mM $MgCl_2$, 5 mM NaCl, 1 mM EDTA) and centrifuged at 10 krcf for 7 minutes to wet the membrane. Then, the filtrate was discarded and the filter was filled with 450 µL of FoB5 buffer and 50 µL of the folded origami sample. The mixture was centrifuged at 7 krcf for 12 minutes. This process was repeated 3 more times. The ACTR-DNA-origami construct was prepared as follows: 25 µL of filter purified DNA origamis bearing the protein docking sequence were mixed with 1 µL of the AF647-labeled ACTR protein at 10 µM. The mixture was heated to 37°C and slowly cooled to 20 °C over the course of 1 hr using a thermocycler. Samples were subjected to another 3 rounds of filter purification as described previously.

##### DNA sequences

All sequences needed to synthesize the DNA origami were described previously. [1] Here we summarize the additional oligonucleotides needed for the experiment.

- 5'-TAATCACTGTTGCCCTGATTAAATACGTTAATAGTGATGTAGGTGGTAGAGGAA-3' staple for binding ACTR in the hotspot
- 5'-TTCCTCTACCACCTACATCAC-3' ssDNA labelled with maleimide used to tag ACTR
- 5'-TAATCACTGTTGCCCTGATTAAATACGTTAATATGTCTGCTACTCTGTCTG-3' staple with docking site labelled with either Cy5B or AlexaFluor 647 at the 3'-end
- 5'-GACAGA-3' short ssDNA labelled with either Dy 751 or ATTO 740 at the 3'-end

The dye labelled oligonucleotides where obtained from biomers.net GmbH, Germany, except for the ATTO 740-labelled strand, which was obtained from Eurofins Genomics GmbH, Germany and the Cy5B labelled strand, which was produced in-house using Cy5B maleimide and C6-Amino-labelled DNA obtained from Eurofins Genomics GmbH, Germany as previously described. [2]

#### 1.2 Nanoantenna preparation

DNA functionalized spherical gold nanoparticles (100 nm diameter) were produced as described previously [1], with slight modifications: 2 mL of a 0.025 mg/mL solution of 100 nm gold nanoparticles were mixed with 20 µL of 10% Tween 20 and 20 µL of potassium phosphate buffer (4:5 mixture of 1 M monobasic and 1 M dibasic potassium phosphate, Sigma Aldrich, USA) as well as 4 nmol of thiol-modified ssDNA (5'-T20-SH-3', Ella Biotech GmbH, Germany) dissolved in 20 µL ultrapure $H_2O$ and 30 µL of TCEP in ultrapure $H_2O$ at 60 mM. The solution was then stirred on a magnetic stirrer at 40 °C for 1 hr. Then, PBS with additional

3.3 M of NaCl was added stepwise to a final concentration of 750 mM NaCl. Then, the particle solution was mixed 1:1 with PBS supplemented with 10 mM NaCl, 2.11 mM P8709 buffer (Sigma Aldrich, USA), 2.89 mM P8584 buffer (Sigma Aldrich, USA), 0.01% Tween 20 and 1 mM EDTA. The solution was centrifuged for 8 min at 2.8 krcf and the supernatant was discarded. This procedure was repeated 5 more times. Then, the supernatant was discarded and the particles were resuspended in 10 mM TRIS, 1 mM EDTA, 750 mM NaCl buffer to an extinction at 550 nm of approximately 0.1 as measured on a UV-VIS spectrometer (NanoDrop2000, Thermo Fisher Scientific, USA). The immobilized DNA origamis were then incubated at RT overnight with this solution to form the nanoantennas.

#### 1.3 Protein expression and labelling

##### Protein expression

For NCBD, a construct with a single cysteine residue and proline residues 20 and 23 replaced by alanine (to suppress kinetic heterogeneity due to peptidyl-prolyl cis/trans isomerization [3]) was generated by site-directed mutagenesis. Furthermore, the expression construct contained an N-terminal His6-tag cleavable with HRV 3C protease (sequence of the cleaved construct: `GPNRSISPSA LQDLLRTLKS ASSAQQQQQV LNILKSNPQL MAAFIKQRTA KYVANQPGMQ C`). NCBD was co-expressed [4] with wild-type ACTR from a pET-47b(+) vector. Expression was carried out in *Escherichia coli* C41(DE3) (Merck). Cells were grown at 37 °C in TYH medium (for 1 L: 20 g tryptone, 10 g yeast extract, 11 g HEPES, 5 g NaCl, 1 g $MgSO_4$, pH 7.3), supplied with 0.5% (w/v) glucose, until they reached an OD600 of 0.8. Then, 1 mM IPTG was added to the culture. Expression continued for 1 h at 37 °C, after which cells were harvested by centrifugation. The harvested cells were lysed by sonication, and the $His_6$-tagged protein was enriched via immobilized metal ion affinity chromatography on Ni-IDA resin (Agarose Bead Technologies). The $His_6$-tag was then cleaved with HRV 3C protease and separated from the protein by another round of IMAC. Finally, NCBD was separated from ACTR and other impurities via reversed-phase HPLC (RP-HPLC) on a C18 column (Reprosil Gold 200, Dr. Maisch, Germany) with a water/0.1% trifluoroacetic acid-acetonitrile gradient. The purified protein was lyophilized, resuspended in buffer, and stored at -80 °C until use. For ACTR, a double-cysteine construct, with one cysteine close to the N-terminus for attachment to the DNA origami, and one close to the C-terminus for the dye, was generated by site-directed mutagenesis. Furthermore, the expression construct contained an N-terminal $His_6$-tag cleavable with HRV 3C protease (sequence of the cleaved construct: `GPCGTQNRPL LRNSLDDLVG PPSNLEGQSD ERALLDQLHT LLSNTDATGL EEIDRALGIP ELVNQGQALE PKQDC`). ACTR was co-expressed with wild-type NCBD from a pET-47b(+) vector and purified as described above.

##### Protein labelling

For NCBD, lyophilized protein was dissolved under nitrogen atmosphere at a concentration of 340 μM in 10 0mM potassium phosphate buffer, pH 7.0, and was labeled for 3 h with a 1.6-fold molar excess of LD750-maleimide (Lumidyne Technologies) to protein. Labeled protein was separated from unlabeled protein with RP-HPLC on a C18 column (Reprosil Gold 200, Dr. Maisch, Germany) with a water/0.1% trifluoroacetic acid-acetonitrile gradient. For ACTR, in a first step, the more reactive cysteine at position 3 was labeled with a 3'-maleimide-functionalized oligodeoxynucleotides (biomers.net GmbH) with the sequence `5'-TTC CTC TAC CAC CTA CAT CAC-3'`-maleimide. Lyophilized protein was dissolved under nitrogen atmosphere to 280 µM in 100 mM potassium phosphate buffer, pH 7.0, and was labeled for 3 h at a 0.5-fold molar ratio of oligodeoxynucleotide to protein. Singly labeled protein was separated from unlabeled and doubly labeled protein with RP-HPLC on a Reprosil-Pur 200 C18-AQ column (Dr. Maisch, Germany), with a water/0.1 M triethylammonium acetate–acetonitrile gradient. In a second step, the cysteine at position 75 was labeled with AlexaFluor647 maleimide. The lyophilized protein/oligodeoxynucleotide construct was dissolved under nitrogen atmosphere to a final concentration of 44 µM in 100 mM potassium phosphate buffer, pH 7.0, and was labeled for 3 h at a 1.5-fold molar ratio of dye to protein. Labeled protein was again separated from unlabeled protein with RP-HPLC on a Reprosil-Pur 200 C18-AQ column, with a water/0.1 M triethylammonium acetate–acetonitrile gradient. The purified constructs were lyophilized, resuspended in buffer, and stored at -80 °C until use.

#### 1.4 Single-molecule surface preparation

Microscopy slides (24 mm $\times$ 60 mm size and 170 µm thickness (Carl Roth GmbH, Germany) were cleaned in an ozonator (PSD-UV4, Novascan Technologies, USA) at 100 °C for 30 mins. Then, 2 CoverWell™ Perfusion Chamber gaskets (9 mm diameter, 0.5 mm deep, Grace Biolabs) were glued on the slides and the glue was strengthened by placing the slide on a 100 °C heating plate for 30 seconds. Surfaces then were cleaned by incubating with 1 M KOH for 10 mins. The chambers then were washed 4 times with PBS. The surfaces then were passivated by incubation with BSA-Biotin (1 mg/mL in 10 mM TRIS, 1 mM EDTA, 50 mM NaCl, Sigma Aldrich, USA) for 10 mins. The chambers were then washed with PBS (3 times). Then, the chambers were incubated with NeutrAvidin™ (0.25 mg/mL in PBS, freshly diluted from a 1 mg/mL stock in ultrapure $H_2O$, Thermo Fisher Scientific, USA) for 10 minutes. After washing with PBS (3 times), chambers were ready for use.

#### 1.5 Confocal microscopy

Data was acquired on a home-built setup based on an Olympus IX-71 microscope body. A LDH-D-C-640 laser (636 nm) in continuous wave mode was focussed to a diffraction-limited spot and sent through a linear polarizer (LPVISE100-A, Thorlabs GmbH) and a quarter-wave plate (AQWP05M-600, Thorlabs GmbH) to obtain circularly polarized light. An oil-immersion objective (UPLSAPO100XO, NA1.40, Olympus Deutschland GmbH) was used to focus the light onto the sample. We used a piezo stage (P-517.3CD, Physik Instrumente GmbH & Co. KG) and a piezo controller (E-727.3CDA, Physik Instrumente GmbH & Co. KG) to scan the sample. The fluorescence was separated from the excitation laser with a dichroic beam splitter (zt488/543/635/730rpc, Chroma Technologies) and focussed on a 50 µm diameter pinhole (ThorlabsGmbH). The fluorescence was then split between the red and infrared channel using a beam splitter (HC BS 749 SP, Chroma). The red fluorescence was distributed to 2

APDs (SPCM-AQRH-14-TR and SPCM-AQR-15, Excelitas Technologies GmbH & Co. KG, Germany) using a nonpolarizing 50:50 beam splitter (CCM1-BS013/M, Thorlabs GmbH). The infrared channel was distributed to 2 APDs (SPCM-AQRH-14-TR and SPCM-AQR-15, Excelitas Technologies GmbH & Co. KG, Germany) using a nonpolarizing 50:50 beam splitter (CCM1-BS014/M, Thorlabs GmbH). Events were registered by a multichannel picosecond event timer (HydraHarp 400, PicoQuant GmbH) in T2 mode and the hardware was controlled using a commercial software (SymPhoTime 64, PicoQuant GmbH) We used a photostabilizing system consisting of enzymatic oxygen removal by glucose oxidase/catalase as well as a reducing/oxidizing system with Trolox/Troloxquinone for all of the experiments. Specifically, the buffer consisted of 50 mM TRIS, 500 mM NaCl, 2 mM Trolox/Troloquinone, 2% (w/v) Glucose, 165 U/mL Glucose oxidase, 2170 U/mL Catalase dissolved in $D_2O$ for all experiments. [5, 6] For the experiments with ATTO 740, the pH value of the buffer was adjusted to 7 before use. We used lower excitation powers (10 nW – 20 nW) to scan the surfaces and find the coordinates of the single molecules. Then we placed the laser focus on these coordinates, checked if the fluorescence transient exhibited binding events and then increased the laser power to 1 µW – 4 µW by removing a neutral density filter in the excitation path.

##### 1.6 Data analysis

Data were analysed using home-written python code with a backend written in C++ using Eigen [7] which is available on Gitlab. [8] For the maximum likelihood analysis of photon-by-photon trajectories we used the method developed by Gopich and Szabo [9] and applied to transition paths by Chung and Eaton. [10–12] Here, the likelihood of detecting a sequence of photons j with colors $c_i$ is given by

$$L_j = \nu_{fin}^{\mathbf{T}} \prod_{i=2}^{N} \left[\mathbf{F}(c_i) \exp(\mathbf{K}\tau_i)\right] \mathbf{F}(c_1)\, \nu_{ini} \tag{1}$$

with $\tau_i$ being the interphoton time and $\mathbf{K}$ being the rate matrix either for the two state model with acceptor blinking:

$$\mathbf{K} = \begin{bmatrix} -k_D - k_d & k_{A,app} & k_b & 0 \\ k_D & -k_{A,app} - k_d & 0 & k_b \\ k_d & 0 & -k_D - k_b & k_{A,app} \\ 0 & k_d & k_D & k_{A,app} - k_b \end{bmatrix} \tag{2}$$

or the model with an intermediate state

$$\mathbf{K} = \begin{bmatrix} -k'_D - k_d & k_i & 0 & k_b & 0 & 0 \\ k'_D & -2k_i - k_d & k'_{A,app} & 0 & k_b & 0 \\ 0 & k_i & -k'_{A,app} - k_d & 0 & 0 & k_b \\ k_d & 0 & 0 & -k'_D - k_b & k_i & 0 \\ 0 & k_d & 0 & k'_D & -2k_i - k_b & k'_{A,app} \\ 0 & 0 & k_d & 0 & k_i & -k'_{A,app} - k_b \end{bmatrix} \tag{3}$$

Here, $k_\text{D}$ is the dissociation rate constant, $k_\text{d}$ and $k_\text{b}$ are the rate constants for the conversion between bright and dark states, $k_\text{A,app}$ is the apparent association rate and $k_\text{i}$ is the rate with which the intermediate state is depopulated. **F** is the photon color matrix and is given by

$$\mathbf{F}_\text{acceptor} = \begin{bmatrix} E_\text{B} & 0 & 0 & 0 \\ 0 & E_\text{U} & 0 & 0 \\ 0 & 0 & E_\text{d} & 0 \\ 0 & 0 & 0 & E_\text{d} \end{bmatrix} \quad \text{and} \quad \mathbf{F}_\text{donor} = I - \mathbf{F}_\text{acceptor} \tag{4}$$

for the two state model and

$$\mathbf{F}_\text{acceptor} = \begin{bmatrix} E_\text{B} & 0 & 0 & 0 & 0 & 0 \\ 0 & E_I & 0 & 0 & 0 & 0 \\ 0 & 0 & E_\text{U} & 0 & 0 & 0 \\ 0 & 0 & 0 & E_\text{d} & 0 & 0 \\ 0 & 0 & 0 & 0 & E_\text{d} & 0 \\ 0 & 0 & 0 & 0 & 0 & E_\text{d} \end{bmatrix} \quad \text{and} \quad \mathbf{F}_\text{donor} = I - \mathbf{F}_\text{acceptor} \tag{5}$$

for the model with an intermediate state. Here, $I$ is the identity matrix, $E_\text{B}$ is the FRET efficiency in the bound state, $E_\text{U}$ is the FRET efficiency in the unbound state and $E_I$ is the FRET efficiency of the intermediate state. $E_\text{d}$ is the FRET efficiency of the dark state, which we set equal to $E_\text{U}$. Because we are analyzing single transitions from the unbound to the bound state, we set $k_\text{A,app} = k_\text{D} = 0.01s^{-1}$, $k'_\text{A,app} = k'_\text{D} = 0.02s^{-1}$ and

$$\nu_\text{ini}^\mathbf{T} = \begin{pmatrix} 0 & p_\text{b} & 0 & 1 - p_\text{b} \end{pmatrix} \quad \text{and} \quad \nu_\text{fin} = \begin{pmatrix} p_\text{b} & 0 & 1 - p_\text{b} & 0 \end{pmatrix} \tag{6}$$

for the two state model and

$$\nu_\text{ini}^\mathbf{T} = \begin{pmatrix} 0 & 0 & p_\text{B} & 0 & 0 & 1 - p_\text{B} \end{pmatrix} \quad \text{and} \quad \nu_\text{fin} = \begin{pmatrix} p_\text{B} & 0 & 0 & 1 - p_\text{B} & 0 & 0 \end{pmatrix} \tag{7}$$

for the model with an intermediate state with $p_\text{b} = k_\text{b}/(k_\text{b} + k_\text{d})$. For the fitting procedure, it is beneficial to define $k = k_\text{b} + k_\text{d}$. We manually picked single transitions and fit them with the two state model to obtain $E_\text{B}$, $E_\text{U}$, $k$ and $p_\text{b}$. We discarded transitions where the fit did not converge (41 and 33 discarded transitions for the experiment with the IDPs and the experiments with the DNA hybridization, respectively) and where $E_\text{B}$ was more than 2 $\sigma$ away from the mean value (7 and 19 discarded transitions). Then we used the obtained fit parameters to calculate $\Delta \ln L$ with $\tau_\text{i} = 1/(2k_\text{i})$ and $E_I$ as the only free parameters. To summarize the results, we normalized the obtained $E_I$ values to the respective $E_\text{U}$ and $E_\text{B}$ values via $E_\text{i, norm} = (E_I - E_\text{U})/(E_\text{B} - E_\text{U})$ for each transition.

### 2 Supplementary notes

#### 2.1 Considerations on the choice of the FRET pair

The most important parameters for the choice of a suitable FRET pair are the absorption spectra of the dyes with respect to the plasmon resonance of the nanoantenna, because the fluorescence enhancement is the region slightly red shifted to the plasmon resonance. The plasmon resonance peak can be approximated by the scattering cross-section of the nanoantenna. As shown previously [13], for dimers of 80 nm silver nanoparticles (AgNPs), the plasmon resonance lies at 500–650 nm while for 100 nm gold nanoparticles (AuNPs) it lies in the region of 600–750 nm. Because of increased scattering of AgNPs, we opted to utilize nanoantennas based on AuNPs, which meant that we had to work with dyes absorbing in the red to near-IR spectral range. We also noticed a negative influence of pulsed excitation on the photostability of the fluorophores which is why we used CW excitation at 640 nm.

The next considerations were possible dim or dark states of the fluorophores. We showed that carbopyronine dyes such as ATTO 647N show pronounced dim state formation in nanoantenna environments. [14] On the other hand, cyanine dyes did not show such behavior: dark state formation, e.g. by trans-cis isomerization was accelerated, but no second intensity state was observed, which makes these class of dyes superior for this application. For this reason, we chose cyanine dyes as donor dyes – either AlexaFluor 647 or Cy5B. [2]

For the acceptor fluorophores, the same considerations apply, but the available options of near-IR absorbing dyes are more limited. They comprise mainly cyanine based dyes such as AlexaFluor 750, Lumidyne (LD) 750 or Dyomics (Dy-)751 and as an alternative ATTO740 (structure unpublished, to the best of our knowledge). We tried most of them with similar success so that the more deciding factor in this regard was the attainable proximity ratio in the bound state, which is likely influenced by Dye-DNA or Dye-Protein interactions and the compatibility with our labelling reaction (ATTO740 is pH sensitive, LD-750 and AlexaFluor 750 were not commercially available on very short oligonucleotides). We note that in all cases, the use of ROXS and oxygen removal leads to sub-microsecond acceptor blinking [15] which we attribute to trans-cis isomerization as well as ROXS-induced radical blinking. This could potentially be overcome by alternative photostabilization methods which rely on efficient direct depopulation of the triplet excited states (e.g. energy transfer mechanisms). [16]

#### 2.2 Note on the comparison of nanoantenna and reference measurements

To assess the increase in maximum photon count rates and total observation times, the fairest comparison would be to use the same FRET pair (AlexaFluor 647 and LD750) measured without the DNA nanoantenna. However, since the photostability of this red/IR dye pair outside of the plasmonic hotspot was very poor (see exemplary time traces in Figure S7), we could only compare our results in the DNA nanoantenna to the previous state of the art, which in this case was the more photostable green/red FRET pair consisting of Cy3B and CF660R. [17]

#### 2.3 Simulations of photon time traces

To test the robustness of the analysis based on transfer efficiencies for the case of varying total photon count rates, we simulated photon time traces with varying total photon count rates and subjected them to the analysis. For the sake of simplifying the procedure, we did not include the possibility of photoblinking in these simulations. Time traces were simulated by drawing random interphoton times from an exponential distribution with the respective count rate and assigning them to the donor or acceptor channel with the relative probability given by the FRET efficiency of the given state. The datasets were then analysed with the same procedure, with the rate matrices $K$ for the two state and the three state model, respectively, given by

$$\mathbf{K} = \begin{bmatrix} -k_{\text{off}} & k_{\text{on}} \\ k_{\text{off}} & -k_{\text{on}} \end{bmatrix} \quad \text{and} \quad \mathbf{K} = \begin{bmatrix} -k_{\text{off}} & k_{\text{i}} & 0 \\ k_{\text{off}} & -2k_{\text{i}} & k_{\text{on}} \\ 0 & k_{\text{i}} & -k_{\text{on}} \end{bmatrix} \tag{8}$$

The results are shown in Supplementary Figures 3 and 4. In conclusion, while the total photon count rate during the transition seems to affect the accuracy of the result and especially the total log likelihood difference, the varying photon count rates themselves seem to have only a minor effect on the extracted values.

### 3 Supplementary tables

**Table S1:** Mean fit results for the two state model. Errors indicate standard errors obtained from each dataset.

| Sample | Parameter | Fit result |
|---|---|---|
| ACTR-NCBD<br>AlexaFluor 647/LD 750<br>(141 transitions) | $E_B$<br>$E_U$<br>$k$<br>$p_b$ | $0.489 \pm 0.012$<br>$0.060 \pm 0.002$<br>$1240 \pm 137$ ms$^{-1}$<br>$0.849 \pm 0.012$ |
| DNA hybridization<br>Cy5B/Dy751<br>(405 transitions) | $E_B$<br>$E_U$<br>$k$<br>$p_b$ | $0.467 \pm 0.004$<br>$0.070 \pm 0.001$<br>$412 \pm 49$ ms$^{-1}$<br>$0.880 \pm 0.005$ |
| DNA hybridization<br>AlexaFluor 647/ATTO740<br>(96 transitions) | $E_B$<br>$E_U$<br>$k$<br>$p_b$ | $0.302 \pm 0.007$<br>$0.047 \pm 0.001$<br>$705 \pm 107$ ms$^{-1}$<br>$0.954 \pm 0.004$ |
| DNA hybridization (mismatch)<br>AlexaFluor 647/ATTO740<br>(204 transitions) | $E_B$<br>$E_U$<br>$k$<br>$p_b$ | $0.272 \pm 0.003$<br>$0.042 \pm 0.001$<br>$674 \pm 57$ ms$^{-1}$<br>$0.945 \pm 0.005$ |

**Table S2:** Fit results for the three state model. Errors are calculated from the diagonal elements of the covariance matrix.

| Sample | Parameter | Fit result |
|---|---|---|
| ACTR-NCBD<br>AlexaFluor 647/LD 750 | $\tau_\mathrm{i}$ | 96 ± 13 μs |
| | $E_\mathrm{i,\,norm}$ | 0.58 ± 0.04 |
| | $\Delta \ln L\,(\tau_\mathrm{i},\ E_\mathrm{i,\,norm})$ | 92.146 |
| DNA hybridization<br>Cy5B/Dy751 | $\tau_\mathrm{i}$ | 17 ± 1 μs |
| | $E_\mathrm{i,\,norm}$ | 0.30 ± 0.02 |
| | $\Delta \ln L\,(\tau_\mathrm{i},\ E_\mathrm{i,\,norm})$ | 247.261 |
| DNA hybridization<br>AlexaFluor 647/ATTO740 | $\tau_\mathrm{i}$ | 20 ± 3 μs |
| | $E_\mathrm{i,\,norm}$ | 0.21 ± 0.02 |
| | $\Delta \ln L\,(\tau_\mathrm{i},\ E_\mathrm{i,\,norm})$ | 67.301 |
| DNA hybridization (mismatch)<br>AlexaFluor 647/ATTO740 | $\tau_\mathrm{i}$ | 41 ± 3 μs |
| | $E_\mathrm{i,\,norm}$ | 0.23 ± 0.01 |
| | $\Delta \ln L\,(\tau_\mathrm{i},\ E_\mathrm{i,\,norm})$ | 278.717 |

**Table S3:** Estimated Förster radii for all FRET pairs used in this study.

| FRET pair | Estimated $R_0$ |
|---|---|
| AlexaFluor 647 – LD750 | 75.7 Å |
| Cy5B – Dy751 | 70.9 Å |
| AlexaFluor 647 – ATTO740 | 65.4 Å |

### 4 Supplementary figures

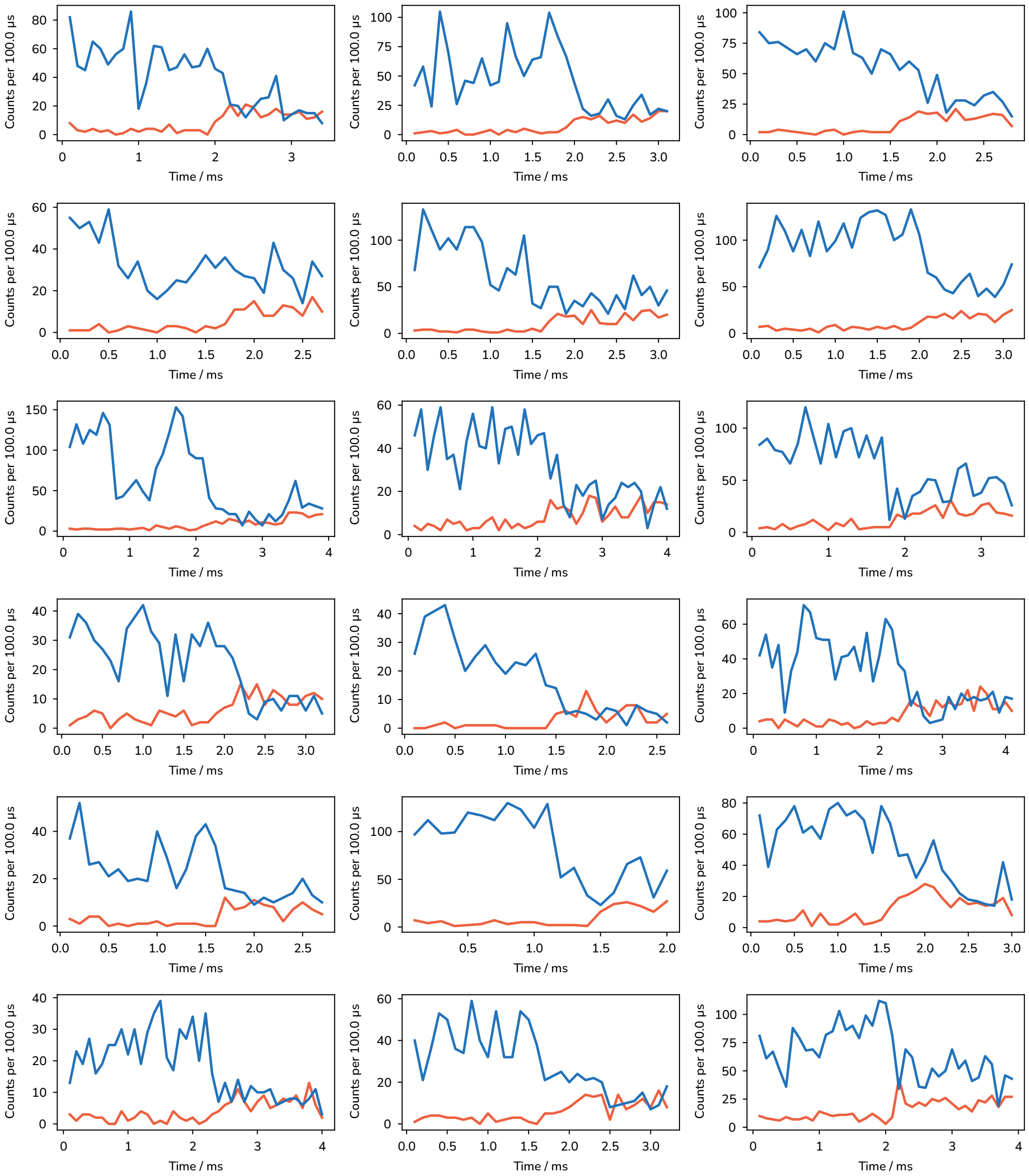

**Figure S1:** Exemplary fluorescence time traces containing transitions for the ACTR-NCBD experiment. Blue: donor fluorescence, red: acceptor fluorescence.

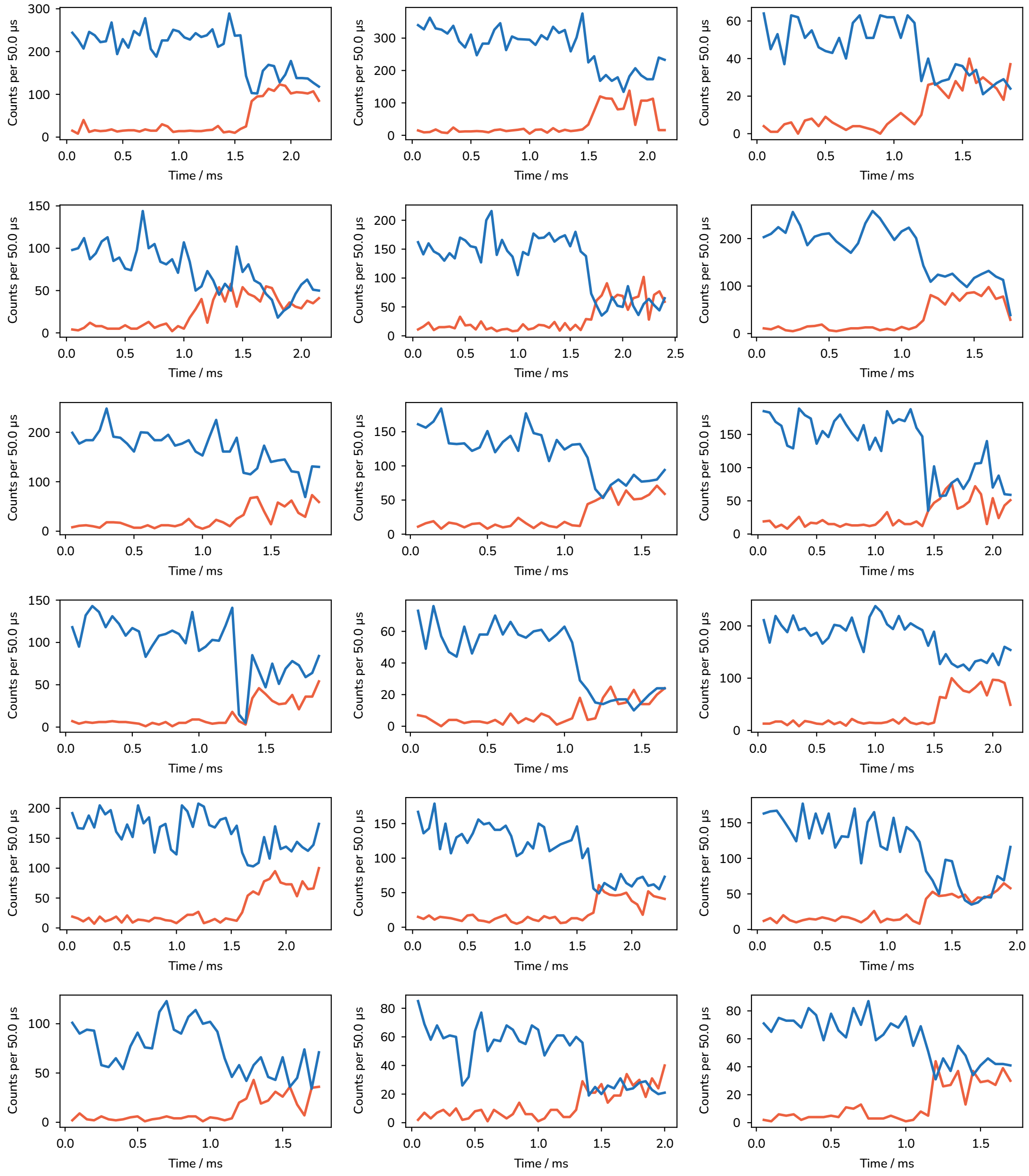

**Figure S2:** Exemplary fluorescence time traces containing transitions for the DNA hybridization experiment. Blue: donor fluorescence, red: acceptor fluorescence.

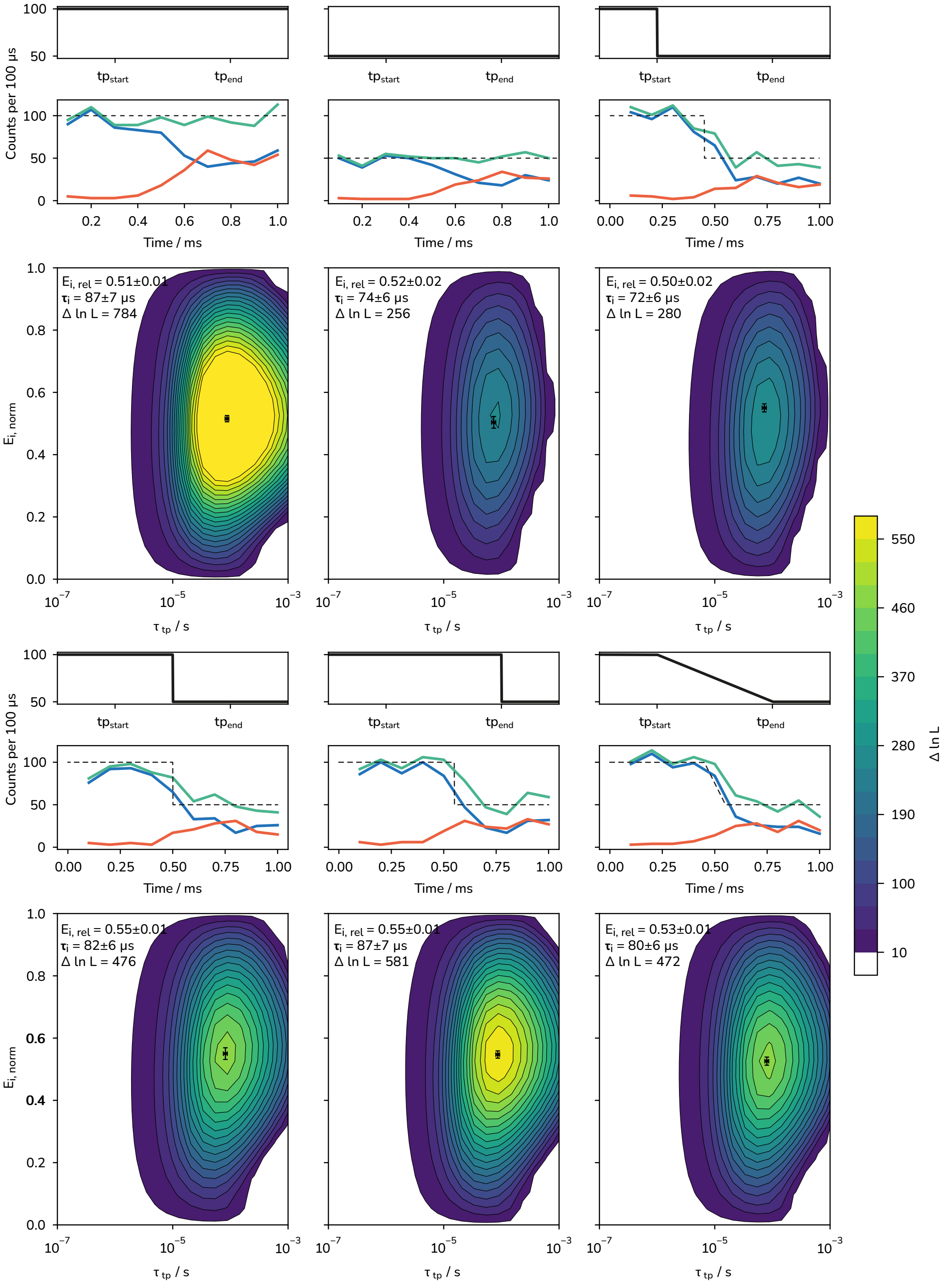

**Figure S3:** Analysis of the influence of varying photon count rates on the robustness of the analysis. For each set, 200 transitions (1 ms windows) were simulated. Input parameters were chosen to ressemble the experiments with the IDPs ($E_\mathrm{B} = 0.5$, $E_\mathrm{i,\,norm} = 0.5$, $E_\mathrm{U} = 0.05$, $\tau_\mathrm{i} = 100$ µs, PCR = 1–0.5 MHz) Blue: donor fluorescence, Red: acceptor fluorescence, Green: sum of both channels, Black: input countrate for the simulation. The most likely values are indicated in the plots. The errors were calculated from the diagonal elements of the covariance matrix. See Supplementary Note 2 for details on the analysis.

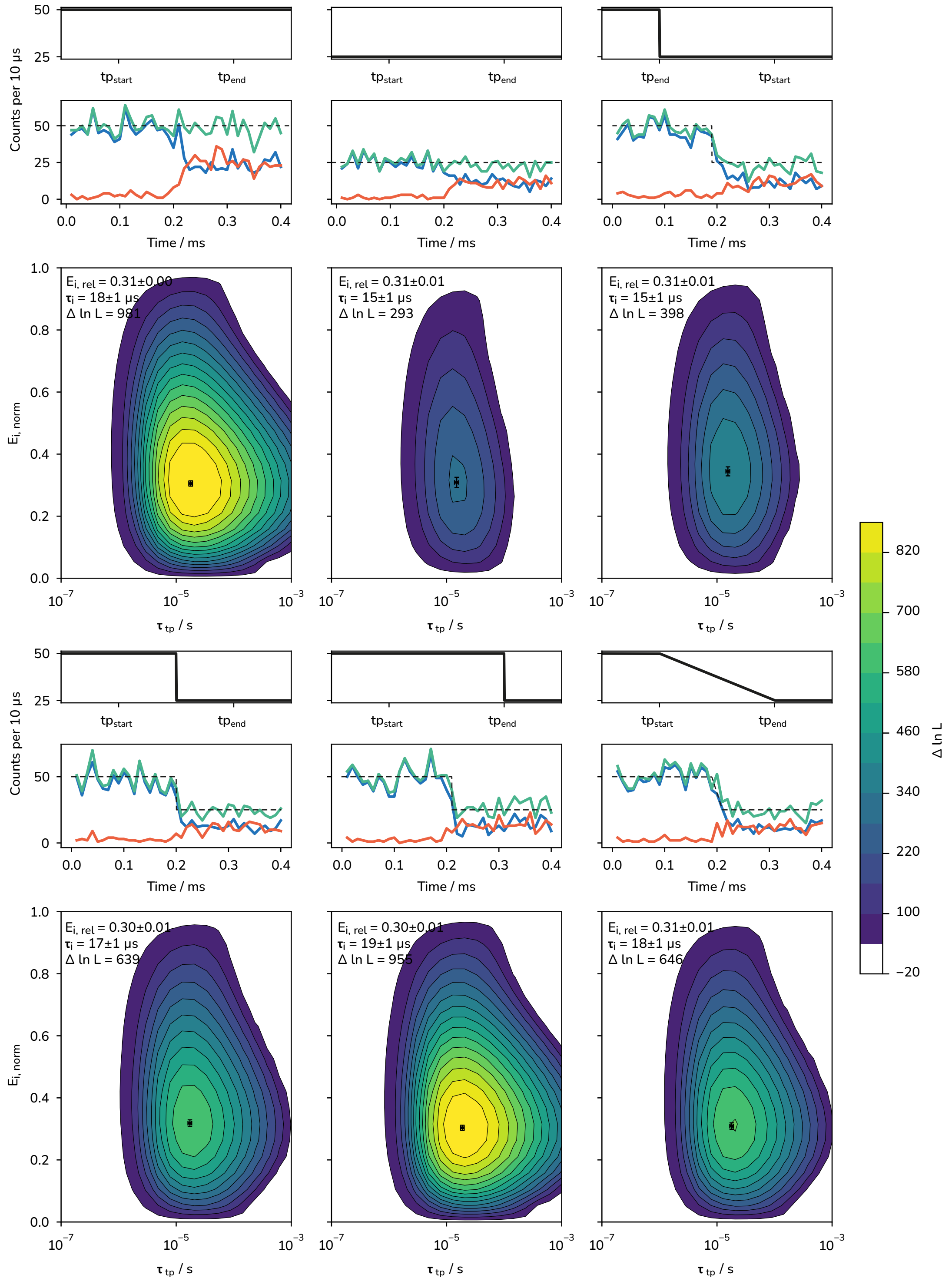

**Figure S4:** Analysis of the influence of varying photon count rates on the robustness of the analysis. For each set, 200 transitions (1 ms windows) were simulated. Input parameters were chosen to ressemble the experiments with the DNA hybridization reaction ($E_\mathrm{B} = 0.5$, $E_\mathrm{i,\,norm} = 0.3$, $E_\mathrm{U} = 0.05$, $\tau_\mathrm{i} = 20$ μs, PCR = 5–2.5 MHz) Blue: donor fluorescence, Red: acceptor fluorescence, Green: sum of both channels, Black: input countrate for the simulation. The most likely values are indicated in the plots. The errors were calculated from the diagonal elements of the covariance matrix. See Supplementary Note 2 for details on the analysis.

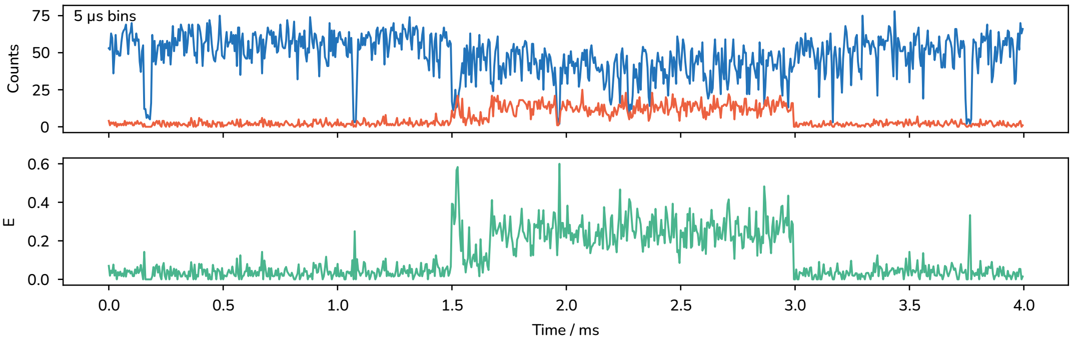

**Figure S5:** Exemplary binding event of the 6 nt long ssDNA to the docking site at 5 μs binning. Blue: donor (AlexaFluor 647) fluorescence, red: acceptor (ATTO740) fluorescence (top). The bottom panel shows the uncorrected FRET efficiency. This snippet contains photon count rates of up to 16.4 MHz (average: 10.8 MHz).

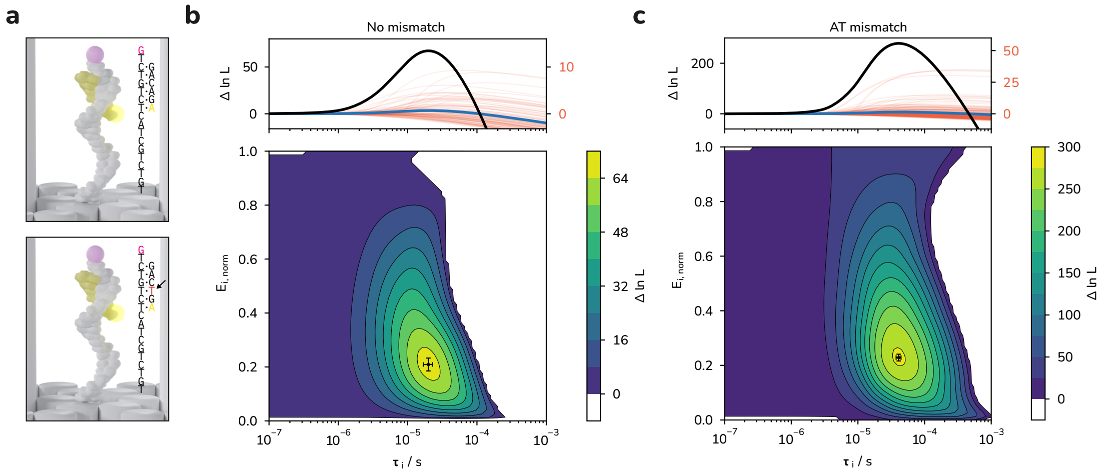

**Figure S6: (a)** Depiction of the position of the AT mismatch in the short ssDNA strand (bottom) as well as the sequence of the perfectly matched staple (top). **(b)** 2D contour plot of the log-likelihood difference $\Delta \ln L$ for the perfectly matched sample with AlexaFluor 647 and ATTO740 summed up over 96 transitions (bottom) as well as a plot showing $\Delta \ln L$ values versus $\tau_i$ for single transitions (red, right scale) and the sum of all transitions (black) as well as their average (blue, right scale) calculated at the most likely value for $E_{i,\,norm}$. The most likely values are $\tau_i = 20 \pm 3$ μs and $E_{i,\,norm} = 0.21 \pm 0.02$. **(c)** Same plots as (b) but for the sample with an AT mismatch in the short ssDNA stand. The most likely values are $\tau_i = 41 \pm 3$ μs and $E_{i,\,norm} = 0.23 \pm 0.01$. The errors were calculated from the diagonal elements of the covariance matrix.

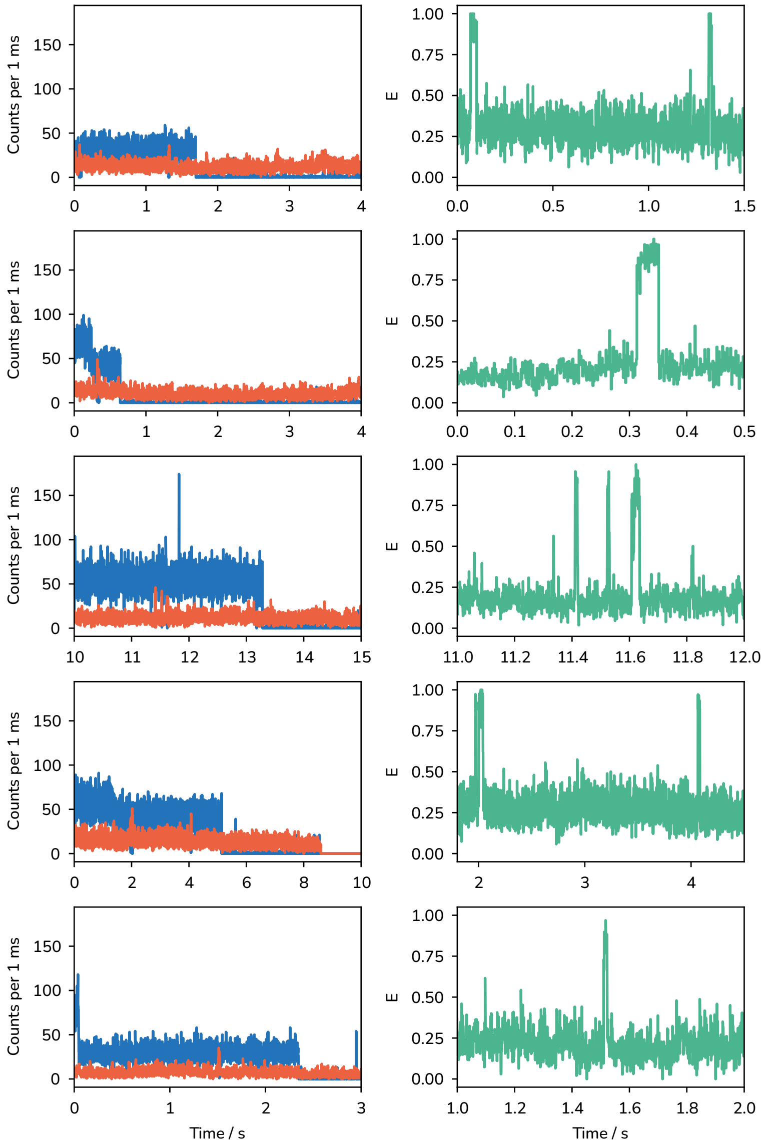

**Figure S7:** Exemplary time traces for the ACTR/NCBD system labelled with AlexaFluor 647 and LD750 immobilized on the DNA origami without attached nanoparticles. Left: donor (blue) and acceptor (red) fluorescence time traces. Right: corresponding zoom-ins on binding events in the uncorrected FRET time traces. Time traces were acquired at 1–2 μW excitation intensity.

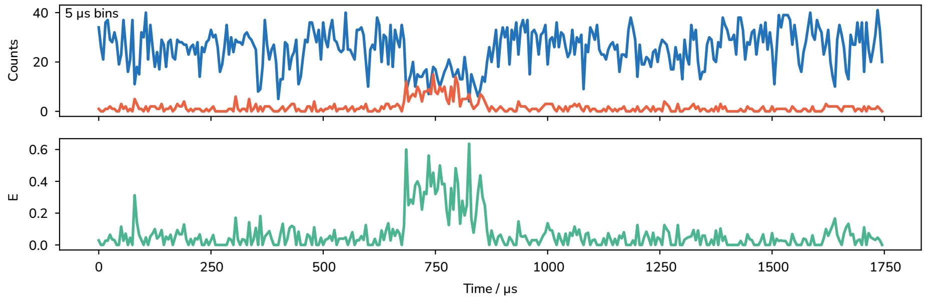

**Figure S8:** Exemplary time trace showing a DNA-DNA hybridization reaction for the sample with an AT-mismatch (estimated $K_d \approx 200$ μM) at 5 μs binning. The concentration of the short DNA strand used for this experiment was 3.6 μM. Blue: donor (AlexaFluor 647) fluorescence, red: acceptor (ATTO740) fluorescence (top). The bottom panel shows the uncorrected FRET efficiency.
